# Assessing the efficiency of catch-up campaigns for introduction of pneumococcal conjugate vaccine; a modelling study based on data from Kilifi, Kenya

**DOI:** 10.1101/118976

**Authors:** Stefan Flasche, John Ojal, Olivier Le Polain de Waroux, Mark Otiende, Katherine L. O’Brien, Moses Kiti, D. James Nokes, W John Edmunds, J. Anthony G. Scott

## Abstract

**Background:** The World Health Organisation recommends the use of catch-up campaigns as part of the introduction of pneumococcal conjugate vaccines (PCVs) to accelerate herd protection and hence PCV impact. The value of a catch-up campaign is a trade-off between the costs of vaccinating additional age groups and the benefit of additional direct and indirect protection. There is a paucity of observational data, particularly from low-middle income countries to quantify the optimal breadth of such catch-up campaigns.

**Methods:** In Kilifi, Kenya PCV10 was introduced in 2011 using the 3-dose EPI infant schedule and a catch-up campaign in children <5 years old. We fitted a transmission dynamic model to detailed local data including nasopharyngeal carriage and invasive pneumococcal disease (IPD) to infer the marginal impact of the PCV catch-up campaign over hypothetical routine cohort vaccination in that setting, and to estimate the likely impact of alternative campaigns and their dose-efficiency.

**Results:** We estimated that, within 10 years of introduction, the catch-up campaign among <5y olds prevents an additional 65 (48 to 84) IPD cases, compared to PCV cohort introduction alone. Vaccination without any catch-up campaign prevented 155 (121 to 193) IPD cases and used 1321 (1058 to 1698) PCV doses per IPD case prevented. In the years after implementation, the PCV programme gradually accrues herd protection and hence its dose-efficiency increases: 10 years after the start of cohort vaccination alone the programme used 910 (732 to 1184) doses per IPD case averted. We estimated that a two-dose catch-up among <1y olds uses an additional 910 (732 to 1184) doses per additional IPD case averted. Furthermore, by extending a single dose catch-up campaign to children 1 to <2y old and subsequently to 2 to <5y olds the campaign uses an additional 412 (296 to 606) and 543 (403 to 763) doses per additional IPD case averted. These results were not sensitive to vaccine coverage, serotype competition, the duration of vaccine protection or the relative protection of infants.

**Conclusions:** We find that catch-up campaigns are a highly dose-efficient way to accelerate population protection against pneumococcal disease.

## Introduction

With the aid of Gavi, the Vaccine Alliance (Gavi), many low income countries, in particular across Africa, have introduced pneumococcal conjugate vaccines (PCVs) into their infant immunisation programmes. However, there remain Gavi countries particularly in south Asia and northern Africa, some with large infant populations, who are yet to follow [1]. Country policy makers, along with global stakeholders, have high interest in achieving optimal health impact from PCV as quickly as possible, however, approaches for achieving maximum and rapid impact have to be weighed against relative cost. In situations where vaccine supply is constrained, as was the case several years ago for PCV, issues of efficiency and equity in vaccine use are also a consideration [2]. The World Health Organisation (WHO) recommends that catch-up campaigns can be used as part of the introduction of PCVs to accelerate the build-up of herd protection and hence PCV impact [3]. However, it is unclear if such catch-up campaigns are an efficient way to use PCV or if the gains from such approach are less than the relative increase in the number of doses required.

The value of a catch-up campaign is assessed by quantifying the trade-off between the costs of vaccinating additional age groups and the benefit of additional direct and indirect protection. However, there are few observational data on the impact of PCV campaigns, particularly from low-and middle income countries (LMICs), to quantify the optimal approach of catch-up campaigns. One of the few well-studied examples of a PCV introduction catch-up campaign in a LMIC occurred in Kilifi, Kenya. The 10- valent pneumococcal non typeable *Haemophilus influenzae* protein D-conjugate vaccine (PCV10) was introduced into the Kenyan routine childhood vaccination programme in early January 2011 using the WHO Expanded Programme on immunization (EPI) schedule of 3 infant doses at 6, 10 and 14 weeks. Additionally, in Kilifi County, at the introduction of the cohort programme a 3 dose catch-up campaign was offered to all infants less than 12 months of age and a 2 dose catch-up to children 12-59 months of age.

We fitted a transmission dynamic model of pneumococcal carriage (a precondition for disease and the source of person to person community transmission) and disease to detailed pre- and post PCV introduction data from Kilifi. We aimed to quantify the marginal impact of the PCV catch-up campaign on carriage and disease in Kilifi over the hypothetical impact of a routine cohort vaccination programme alone in that setting. The model also allowed estimation of the likely impact and efficiency of alternative catch-up campaigns.

## Methods

### Data

#### Study population and mixing patterns

Kilifi County is a mainly rural area at the Indian Ocean coast of Kenya. The Kilifi Health and Demographic Surveillance System (KHDSS) was established in 2000. Approximately 260,000 people reside in the KHDSS area and 60% are younger than 20 years of age [4]. Within the KHDSS numerous studies regarding pneumococcus and it’s health effects have been conducted that informed this work (Table 1). The demographic structure of the model is based on 2009 mid-year population census estimates and assumes no demographic changes with time. To adjust for changes in the population age distribution we used respective annual mid-year population estimates to calculate the invasive pneumococcal disease (IPD) incidence rates. A cross sectional prospective diary-based contact survey was conducted in the northern part of KHDSS in 2009 [5,6]. In total 623 randomly selected participants of all ages produced 568 completed diaries in which they reported their contacts during 24 hours and reported 27,395 physical (i.e. skin to skin) contacts with 10,042 unique individuals. This information was used as a proxy for transmission of pneumococcal carriage [7,8]. Standard methods were used to calculate the WAIFW (Who Acquires Infection From Whom) mixing matrix for age groups <1y, 1-5y, 6-15y, 16-19y, 20-49y and older than 50 years for Kilifi HDSS [5,8–10].

**Table 1:**
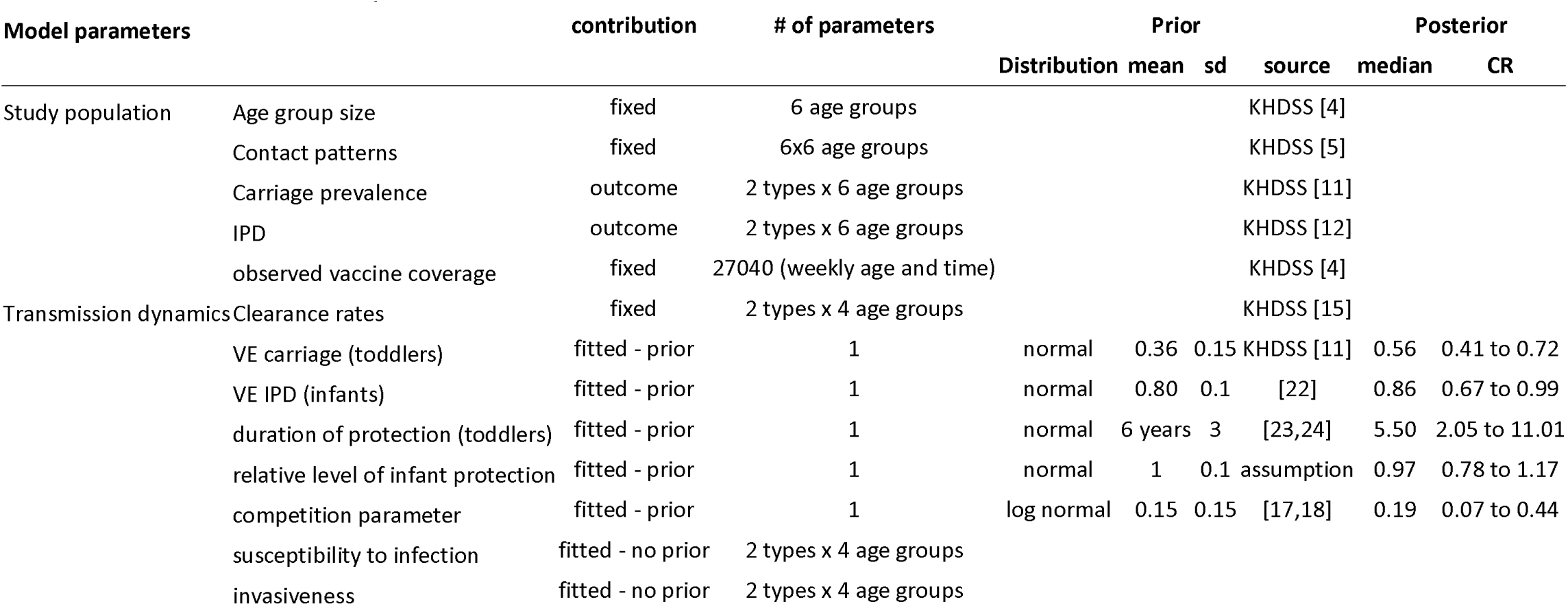
Overview of model parameters.

#### Pneumococcal carriage and IPD

The model was fitted to vaccine type and non-vaccine type carriage prevalence and IPD incidence between 2009 and 2015. During that period annual cross sectional carriage surveys were conducted in Kilifi HDSS [11]. In each study a nasopharyngeal swab was collected from more than 500 randomly selected individuals of all ages. Surveillance with passive case finding for IPD was introduced at Kilifi County Hospital in 1998 for children and in 2007 for adults. Among the residents of Kilifi HDSS, 30 to 70 cases of IPD have been reported annually [12]. Much of that variation is due to changes in disease caused by serotype 1, which has been reported to be unstable in various settings before [13,14].

#### Duration of carriage

We used average age-specific pneumococcal colonization clearance rates estimated from a longitudinal carriage survey in Kilifi HDSS [15] and reported for the age groups <22 months, 22-40 months and 41-59 months. Based on other studies [16], we assumed that clearance rates in individuals older than 5 years of age were 60% higher than in children of age 2-4 years.

#### Serotype competition

As the competition parameter, which determines the proportion by which the likelihood of acquisition is reduced by heterologous carriage, based on local data was only estimated serotype specific [15] rather than for pooled vaccine type and non-vaccine type groups we used a log-normal prior distribution with a median of 0.11 based on estimates from other settings [17–19].

#### Vaccine coverage

As part of KHDSS, electronic individual-based records of the delivery of vaccines are routinely collected at vaccine clinics [20]. We calculated weekly estimates of PCV coverage for the two years after PCV introduction; each stratified by weekly age cohorts from newborns up to 5 years of age. Two such coverage estimates were calcualted: vaccine coverage of at least two doses of PCV administered before the age of 1 year, which was deemed “infant protection”, and vaccine coverage of at least one dose of PCV administered after the age of 1 year, deemed “toddler protection”. The choice of at least 2 doses for infants and at least one dose for toddlers was chosen on the basis of observed coverage rates. For calculation of the number of doses used we assumed that vaccinated infants within the routine program received 3 doses, infants vaccinated as part of the catch-up received 2 doses and toddlers received 1 dose. Data for vaccination rates were available only through late 2012; we extrapolated those rates forward in time by assuming the coverage rates as of later 2012 to continue for the rest of the study period.

#### Vaccine efficacy

The efficacy against VT nasopharyngeal carriage of a single dose of PCV10 administered to children 12-59 months old has been estimated in a randomised controlled trial in Kenya at 36% (95% CI:-1 to 60) [21]. We further assumed that vaccine efficacy against VT IPD of a complete primary series was 80% based on a meta-analysis for PCVs for infants elsewhere [22]. These two estimates of vaccine efficacy against VT carriage and VT IPD were used as priors in the fitting process. Those who were vaccinated in infancy, i.e. before one year of age, may have different vaccine efficacy against acquisition of colonisation, progression to invasive disease and the duration of protection than vaccinated toddlers. Hence we allowed for the model to estimate these parameters for infants as a common proportion of that of toddlers, under the null-hypothesis that no difference exists.

#### Duration of vaccine protection

As estimates of the duration of protection from PCV were not available from studies within the KHDSS we used estimates derived from external studies. Hence our prior on the duration of protection against carriage and disease is centered around 6 years [16,23,24].

### Model

We used a Susceptible, Infected & Infectious, Susceptible – type model of the transmission of grouped vaccine and non-vaccine pneumococcal serotypes as described previously [16,17]. The group of vaccine serotypes consisted of all pneumococcal serotypes targeted by PCV10, i.e. serotypes 1, 4, 5, 6B, 7F, 9V, 14, 18C, 19F and 23F. Individuals were grouped into compartments by their age (weekly age groups until 5 years of age and yearly age groups thereafter), their infection status (either susceptible, infected with a vaccine serotype, a non-vaccine serotype or both at the same time) and by their vaccination status (unprotected, infant protection, toddler protection).

Adaptive Markov chain Monte Carlo methods were used to fit the model to the observed data (Figure 1) [25]. A Poisson likelihood was used for IPD and a multinomial likelihood was used for carriage prevalence. We used a Metropolis Hastings algorithm to create samples from the posterior parameter distributions. Prior information was used according to their availability as described earlier (Table 1 and Figure 1).

**Figure 1:**
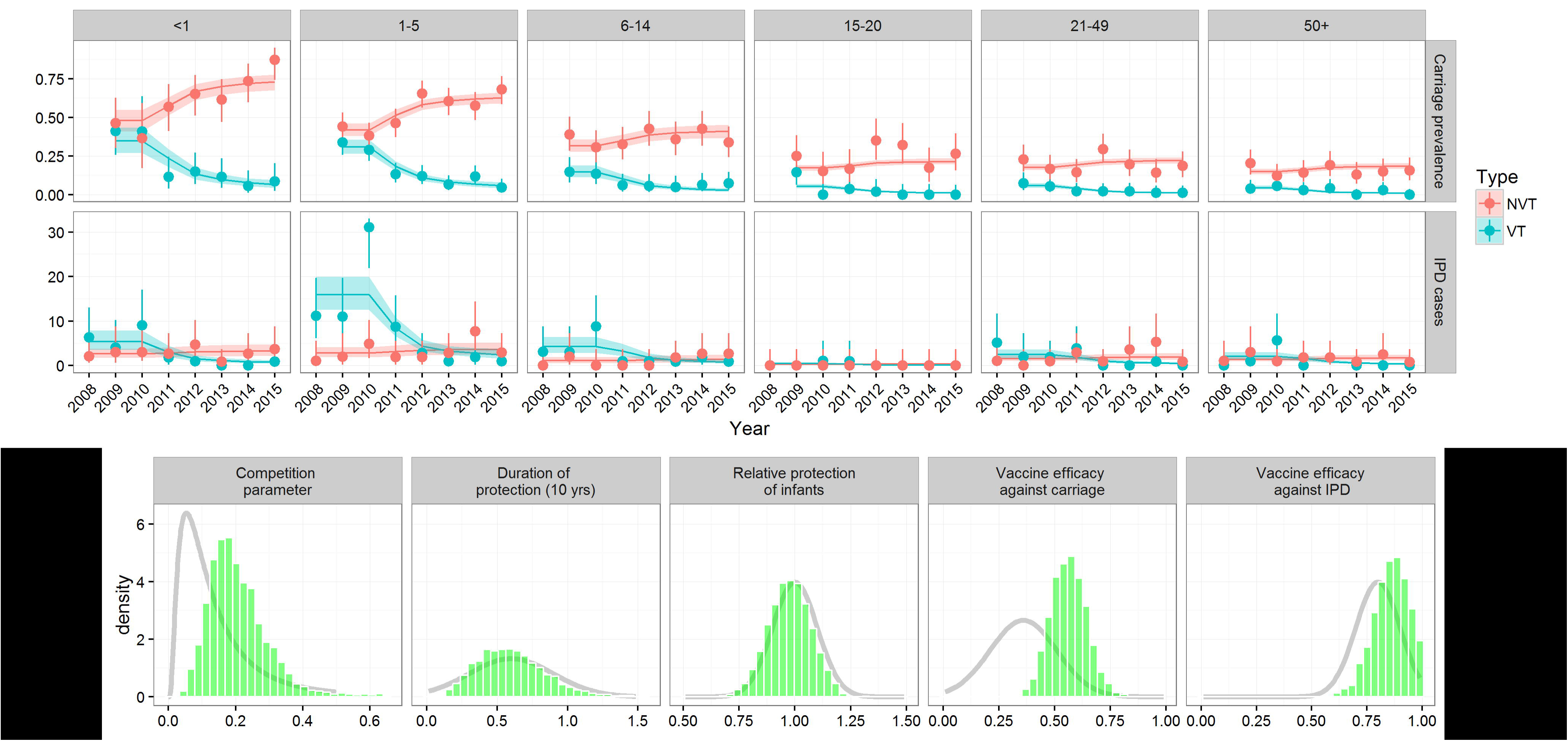
Model fit to carriage prevalence and IPD incidence (upper panel) and prior and posterior parameter estimates (lower panel). Points with 95% confidence bounds represents data and lines with ribbons represent median model estimates with 95% credible intervals. In the lower panel the grey line indicates the prior density distribution and the bars the posterior sample.

### Vaccination scenarios

After fitting the model parameters to match the observed rates of pneumococcal carriage and disease in Kilifi HDSS we created multiple hypothetical PCV introduction scenarios to determine what would have happened if PCV10 had been introduced using different catch-up strategies. For this we define 3 alternative vaccination scenarios which assume that administration of vaccines followed exactly the vaccine uptake that was observed in Kilifi except for the respectively excluded parts of the catch-up programmes. These scenarios were:

1. U5 catch-up (observation and extrapolation) - Vaccination according to observed vaccine coverage in Kilifi HDSS (i.e. all children under 5 years of age).
2. U2 catch-up (hypothetical) - Vaccination according to observed vaccine coverage in Kilifi HDSS for all children under 2 years of age.
3. U1 catch-up (hypothetical) - Vaccination according to observed vaccine coverage in Kilifi HDSS for all children under 1 year of age.
4. Cohort introduction (hypothetical) - Vaccination according to observed vaccine coverage in Kilifi HDSS only for those children eligible for vaccination through cohort introduction.

### Sensitivity analysis

We studied how competition, the duration of protection, the relative protection of infants if compared to toddlers and the vaccine efficacy against carriage and IPD within the range of their posterior distribution impacted on our main outcome; i.e. the number of vaccine doses needed to prevent a case of IPD (NVN) and the ratio of NVN’s of the considered introduction strategies. We used a multivariable linear regression model on the centred posterior samples and report the 95% credible interval limits of the joint distribution of the respective parameter posterior and the model coefficient as a measure of the sensitivity of the NVN ratio to the considered parameters.

We separately assessed the sensitivity of our finding to variable coverage levels in a univariate sensitivity analysis. Rather than the observed coverage levels in Kilifi, for this we assumed that protection through routine immunization as well as the catch-up campaign achieved either 80%, 60% or 40% coverage.

## Results

The introduction of PCV10 together with a catch-up campaign in under 5 year old children was predicted to prevent 220 (172 to 270) cases of IPD in Kilifi within the first 10 years after the start of the vaccination programme. Once the full direct and indirect effects of the programme are established the vaccination programme was predicted to avert 23 (17 to 28) cases of IPD annually (Figure 2a); the majority of those among children (Figure 1).

**Figure 2:**
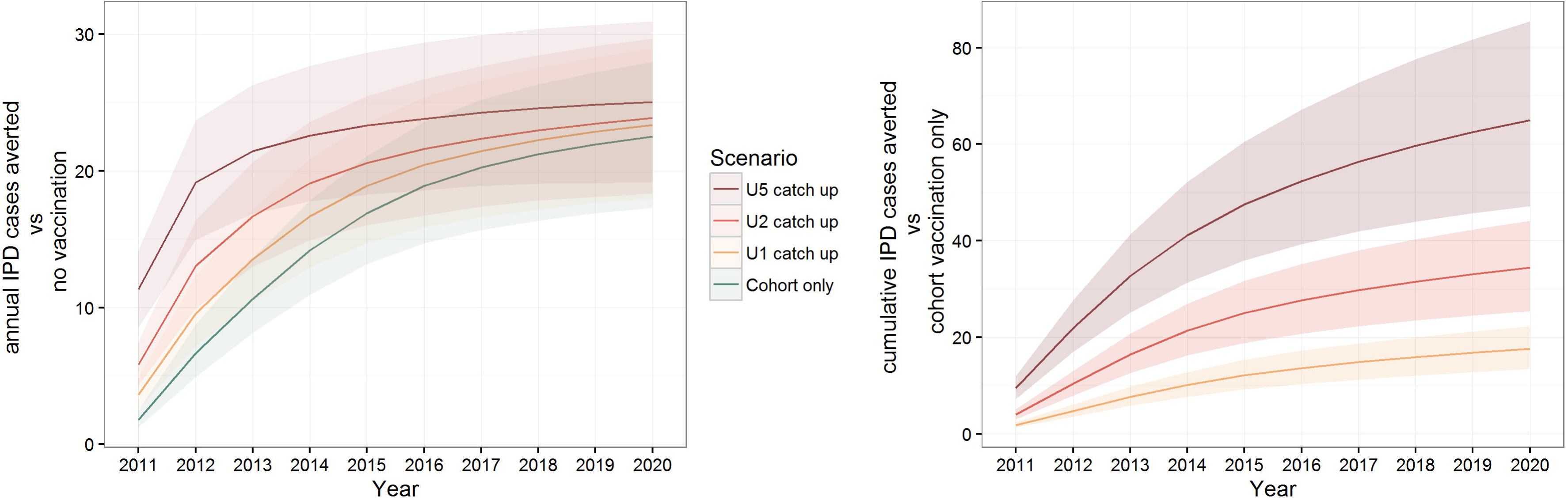
The predicted number of cases averted by PCV10 vaccination in Kilifi if introduced with a catch-up campaign in less than 5, 2 or 1 year olds and without catch-up campaign. Lines represent median estimates and ribbons 95% credible intervals.

The catch-up campaign among children up to 5 years of age was estimated to accelerate direct and indirect effects of PCV. By doing so the Kilifi programme was estimated to prevent an additional 65 (48 to 84) cases of IPD (Figure 2b) over 10 years in the overall population, if compared to a cohort introduction without a catch-up campaign. A catch-up programme confined to children less than 2 years or less than 1 year of age was estimated to prevent 34 (26 to 43) or 18 (14 to 22) IPD cases, respectively, in comparison to cohort introduction alone. The majority of cases averted by the catch-up campaigns would have occurred within the first 6 to 8 years after the start of vaccination (Figure 2a).

Within the first 10 years of the PCV infant programme in Kilifi about 205,000 doses of vaccine were predicted to be used as part of the routine immunisation schedule. The under 5 catch-up campaign required 17,000 additional doses of vaccine (Figure 3a). We estimated that, in the 10 years following introduction of PCV10 in Kilifi, routine vaccination without any catch-up campaigns would use 1321 (1058 to 1698) doses of PCV for each case of IPD averted. As herd protection gradually develops this program gains efficiency in the first years after introduction; that is, the annual number of cases prevented increases while the number of vaccinated individuals remains similar (Figure 2a). By the 10^th^ year after the start of cohort vaccination without a catch-up campaign we estimated that routine vaccination uses 910 (732 to 1184) doses of PCV per IPD case averted.

**Figure 3:**
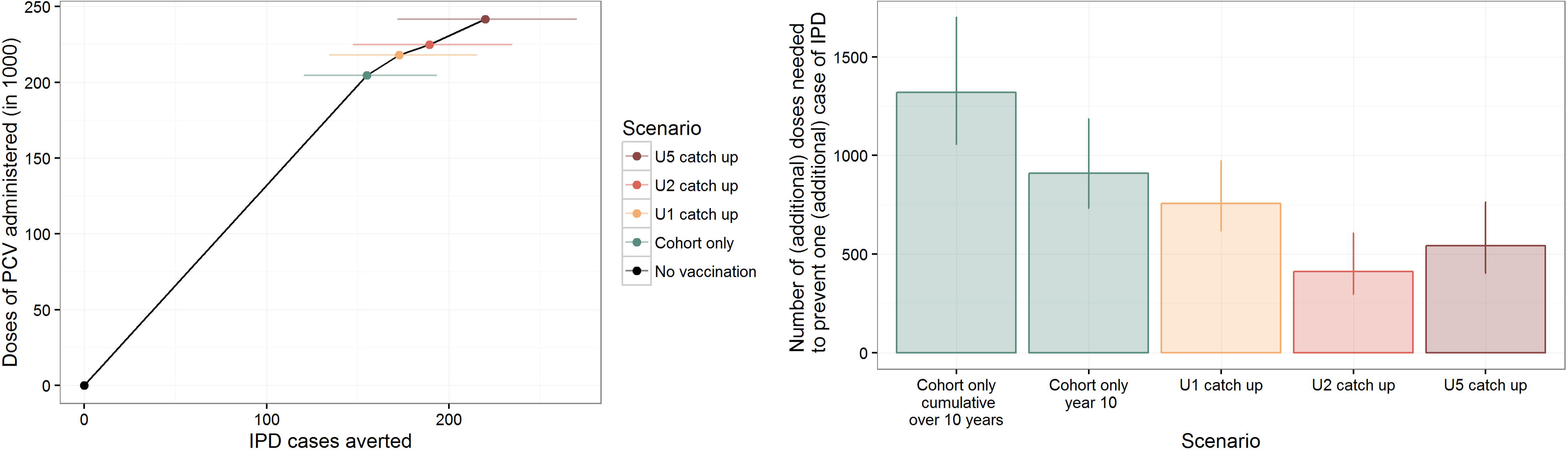
The predicted number of IPD cases averted by PCV10 vaccination in Kilifi in respect to the number of doses administered. In the dose-efficacy plane (left panel) the aggregated dose-efficiency of the alternative introduction strategies within 10 years after the start of vaccination is shown. Coloured dots and lines represent medians and 95% credible intervals (the number of doses administered is fixed as taken from the health register). In the right panel the (incremental) number of doses needed to prevent one (additional) case of IPD. Figures for cohort vaccination alone and cohort vaccination in year 10 are presented as absolute values, the catch-up scenarios are presented as incremental values over the next smaller campaign.

The number of vaccine doses needed to prevent a case of IPD under the four scenarios is shown in Table 2. Extending catch-up PCV immunization to children in the second year of life is the most efficient additional use of PCV but any catch-up campaigns is more efficient than routine birth cohort immunization. The most efficient introduction strategy for PCV is introduction alongside an under 5 year old catch-up.

**Table 2:** The impact and efficiency of alternative introduction strategies. The number of vaccine doses needed to prevent a case of IPD (NVN) is used as a measure of efficiency. Incremental NNV refers to the additional number of doses needed to prevent one additional cases of IPD in respect to cohort introduction with the next smaller catch-up.

| Introduction of PCV<br>via | IPD averted after 10<br>years | Doses<br>administered | Incremental<br>NVN | NVN |
| --- | --- | --- | --- | --- |
| Cohort only | 155 (121 to 193) | 204,671 | 1321 (1058 to 1698) | 1321 (1058 to 1698) |
| + U1 catch-up | 173 (134 to 216) | 218,089 | 757 (618 to 973) | 1263 (1012 to 1623) |
| + U2 catch-up | 189 (147 to 235) | 224,952 | 412 (296 to 606) | 1188 (958 to 1527) |
| + U5 catch-up | 220 (172 to 270) | 241,546 | 543 (403 to 763) | 1098 (894 to 1405) |

Our results were not sensitive to variations in vaccine coverage (Figure S2 and Table S1) or competition, the duration of protection, the relative protection of infants by PCV as compared to toddlers and the vaccine efficacy against carriage and IPD (Figure S1).

## Discussion

In many high-income countries PCVs have been introduced with the help of catch-up campaigns to accelerate the direct and indirect protection that is offered to the community [26–28]. We used extensive data from the Kilifi HDSS, a well-studied mix of rural and urban Kenyan communities representing a typical low income setting, to estimate the incremental effects that different catch-up campaigns are likely to have over routine vaccination and, therefore, whether PCV catch-up campaigns are an efficient use of PCV supply. We found that rapidly increasing the protection in the community via catch-up not only reduces cases of IPD by direct protection of older children but also reduces the burden of IPD in the whole childhood population by developing herd protection more rapidly. Any of the three catch-up programs considered in the analysis were estimated to use fewer vaccine doses to prevent a case of IPD than cohort introduction during the first 10 years; the catch-up schedules were more efficient than routine cohort vaccination programme alone even after full herd effects are in place, in the 10^th^ year of the programme. While the catch-up doses given to one year olds were estimated to be the most efficient ones, we find that cohort introduction alongside a catch-up campaign in under 5 year old children was the most efficient introduction strategy overall.

Data on the observed impact of PCV catch-up campaigns is sparse and mostly circumstantial. Catch-up campaigns of different sizes have been used for introduction of PCVs into countries including the UK, USA, Israel, Brazil and Kenya, however, a head to head comparison with cohort introductions that would allow an evaluation of the additional impact of the catch-up is challenging because of the dissimilarity of the underlying population and other factors including vaccine coverage, intensity of pneumococcal transmission, differences in demographic structure and population mixing, serotype distribution and prevalence of epidemiological risk factors such as HIV infection.

As well as extending direct protection to vulnerable older children, catch-up campaigns also rapidly increase the proportion of individuals in the transmitting population who are protected against VT acquisition and hence onward transmission. This indirect effect is non-linear, preventing a high number of infections for each increment in vaccine coverage when that coverage is low but suffering from a saturation effect for higher coverage levels. As a result, predictions of the optimal extent of catch-up campaigns need to account for these non-linear effects; i.e. incorporate transmission dynamics.

Most of our posterior estimates that had an informative prior were similar to that prior, showing that in most instances the model is able to match the data well using the pre-specified parameter space. The notable exception was the vaccine efficacy against carriage in toddlers. While the model was unable to replicate the observed steep decrease in VT prevalence following vaccination using the mean prior estimate of 36% efficacy the posterior suggests a mean efficacy of 55% which has been observed in other sites [23] and falls well into the range of the prior estimate.

We have restricted our analyses to catch-up campaigns that targeted age groups under five years of age as those were deemed feasible both from a programmatic and a supply point of view. However, including older children may well be efficient in particular in settings where older children contribute substantially to the transmission of pneumococci. Also, we have not considered programmatic issues associated with implementation of catch-up campaigns. Due to the immense additional burden on available staff catch-up campaigns can disrupt routine immunization services. Furthermore, we have studied the most efficient use of PCV supply but not the cost-effectiveness or affordability of catch-up campaigns for PCV introduction. One of the major differences in a cost-effectiveness analysis is that it takes into account the higher delivery costs of vaccine through a supplementary immunization activity. Assuming that doses delivered as part of a PCV catch-up campaign were up 75% more expensive than doses delivered through the routine epi schedule, however, did not qualitatively change our findings on the superior efficiency of catch-up programs.

We did not account for population growth in our model which may impact the transmission dynamics in the post vaccination era and hence on our findings. However, modelling work predicting the impact of PCV10 in Kilifi from pre-vaccination data has shown that accounting for population growth in Kilifi is unlikely to qualitative change the prediction but only slightly reduces the long term impact of vaccination on IPD [29]. As the impact of a catch-up campaign is mostly visible within a few years after vaccination it is likely largely unaffected by long term changes in demographics. Hence, accounting for population growth is likely to further favor the use of catch-up campaigns for introduction of PCV. Other models have taken into account more of the diversity of pneumococcal serotypes by either modelling them individually or by using finer grouping [19,29–31]. Despite the considerable heterogeneity of serotypes in regards to their ecology within both our VT and NVT group our model captures the post-vaccination dynamics well. The impact of catch-up campaigns largely concerns the acceleration of long term impact of the programme and hence nuances in the dynamics of specific serotypes are unlikely to qualitatively change our findings.

The generalisability of our results beyond Kilifi HDSS is dependent on a number of factors. We show in a sensitivity analysis the robustness of our findings to vaccine coverage, vaccine efficacy against the carriage and IPD, the ratio of toddler to infant protection, the duration of vaccine induced protection and the between-serotype group competition (see Appendix Figure S1 and S2). However, other factors that could not be systematically assessed in this analysis include transmission intensity and serotype distribution. In settings with higher transmission intensity, birth cohort PCV introduction likely takes longer to establish full herd effects. As a result, catch-up campaigns that accelerate the build-up of herd protection have the potential to prevent more IPD cases and hence be even more efficient strategy for PCV use. Furthermore, we did not account for potential cross-reactivity of PCV10 against serotypes 6A and 19A which has been reported previously [32,33]. However, both 6A and 19A carriage prevalence increased in the post vaccination era in Kilifi [11].

We assumed that two PCV doses in infancy, given as part of the routine EPI schedule are similarly efficacious at preventing VT carriage and disease as a single catch-up dose in toddlers and young children. Fitting to the data from Kilifi our model did not reject this hypothesis. While our results are robust to factors including this differences in relative protection in infants and toddlers and variable vaccine coverage (proportion of protected infants and toddlers) the number of doses that are administered to establish protection could have a larger impact. Twelve months after the introduction of PCV in Kilifi 76% of infants eligible for 3 doses of PCV aged less than one year had received at least two doses of PCV and 62% of children 1 to 4 years old had received at least 1 dose. We have chosen the dosing of catch-up campaigns to align with what was rolled out in Kilifi, however, other dosing regimens have been used [34], notably South Africa with two doses in infancy followed by a booster dose at 9 months of age [35]. In our analysis we assume for simplicity that all children receive the exact number of doses that in this analysis was deemed sufficient to induce protection. Drop-out rates in Kilifi are relatively low, e.g. more than 97% of infants who received one dose go on to receive a second dose before one year of age, but including drop-outs in the analysis would further decrease the efficiency of the cohort introduction in comparison to the catch-up campaigns. To define protection in our model we used 2 doses in infancy and 1 dose for catch-up campaigns as a protective schedule but assumed that vaccinated children would eventually receive 3 doses as part of the routine schedule or alternatively 2 or 1 dose if part of the catch-up campaign in <1 year old or older children respectively. Assuming instead that the children protected through routine immunization and catch-up campaigns had received 2 doses and 1 doses respectively did not qualitatively change the results.

## Conclusion

Pneumococcal conjugate vaccines are among the most expensive vaccines currently available and make up more than 30% of the annual budget of Gavi. Proposed ways to use PCVs more efficiently include a potential reduction in the number of infant doses if herd effects have been established [36] or a dilution of the current formulation. We show here that catch-up campaigns present an important, readily available tool, which can increase the efficiency of PCVs impact on disease at introduction. For countries yet to introduce, or potentially also for countries with lagging coverage, strategies that include catchup campaigns warrant serious consideration.

## Declarations

### Availability of data and materials

The majority of datasets used and/or analysed during the current study are available from the indicated published resources. The remaining data, including model code is available from the corresponding author on reasonable request.

### Authors’ contributions

JAS and WJE designed the study. SF conducted the analyses, interpreted the results and wrote the first draft of the manuscript. JO, OLPW, MO, KLOB, MK,DJN, WJE and JAS adviced on methodology and / or data interpretation. All authors have contributed to the writing of the manuscript. All authors have read and approved the final manuscript.

## Acknowledgements

We thank the study participants from the Kilifi HDSS, the Ministry of Health District Health Management Team in Kilifi County, and the dedicated team of fieldworkers, data managers, and laboratory scientists who worked on the various studies included in this analysis.

